# Cardiac Sex Differences are Established Prior to Gonad Formation

**DOI:** 10.1101/2020.09.29.319194

**Authors:** Wei Shi, Xinlei Sheng, Kerry M. Dorr, Josiah E. Hutton, Haley A. Davies, Tia D. Andrade, Todd M. Greco, Yutaka Hashimoto, Joel D. Federspiel, Zachary L. Robbe, Xuqi Chen, Arthur P. Arnold, Ileana M. Cristea, Frank L. Conlon

**Author notes:** Correspondence (I.M.C.), (F.L.C.). These authors contributed equally (Wei Shi, Xinlei Sheng).

## Abstract

Male and female disease states differ in their prevalence, treatment responses, and survival rates. In cardiac disease, women almost uniformly fare far worse than men. Though sex plays a critical role in cardiac disease, the mechanisms underlying sex differences in cardiac homeostasis and disease remain unexplained. Here, in adult and embryonic hearts we reveal sex-specific transcriptomes and proteomes and show that cardiac sex differences are predominately accounted for by post-transcriptional mechanisms. We found differential expression of male-female proteins in the cardiomyocytes. Using a quantitative proteomics-based approach, we characterized differential sex-specific enriched cardiac proteins, protein complexes, and biological sex processes in the context of global genetic diversity of the Collaborative Cross, an established surrogate for human diversity. We also found that sex differences in cardiac protein expression are established by both hormonal and sex chromosomal mechanisms. We have demonstrated the onset of sex-biased protein expression and discovered that sex disparities in heart tissue occur at the earliest stages of heart development at a period that preceeds mammalian gonadal development. Collectively, these findings may explain why congenital heart disease, a leading cause of death whose origin is often developmental, is sex biased. Our results reveal molecular foundations for differences in cardiac tissue that underlie sex disparities in health, disease, and treatment outcomes.

## INTRODUCTION

Though sex plays a critical role in cardiac disease, the mechanisms underlying sex differences in cardiac homeostasis and disease remain unexplained. Sex disparities exist in the anatomy and physiology of the heart as well as in the preponderance of specific types of heart disease (Chester et al., 2018; Chlebowski et al., 2017; Dent et al., 2011; Garcia et al., 2016; Haberer and Silversides, 2019; Mosca et al., 2007; Prabhavathi et al., 2014; Whayne and Mukherjee, 2015; Wu et al., 2019). For example, females have a higher rate of atrial septal defects, while males exhibit a higher rate of aortic arch abnormalities (Arnold et al., 2001; Giannakoulas et al., 2017; Maric-Bilkan et al., 2016). Dissimilarities are accentuated in patient outcomes, where women almost uniformly fare far worse than men (Chester et al., 2018; Chlebowski et al., 2017; Dent et al., 2011; Garcia et al., 2016; Haberer and Silversides, 2019; Prabhavathi et al., 2014; Whayne and Mukherjee, 2015; Wu et al., 2019). According to a report published in 2007, one woman dies of cardiovascular disease in the United States every minute (Mosca et al., 2011b).

Clinical studies have implicated menopause as an influencing factor in differing patient outcomes. Women have a lower incidence of cardiac disease until they reach menopause, but post-menopausal women have a rate almost equivalent to males (Regitz-Zagrosek and Seeland, 2011). This observation has led investigators to believe that estrogens plays a protective role (Blenck et al., 2016; Kararigas et al., 2014; Regitz-Zagrosek and Kararigas, 2017; Ventura-Clapier et al., 2017; Wu et al., 2019). However, recent studies have demonstrated that pre-, peri- or post-menopausal hormonal replacement therapies have little effect on the incidence of heart disease (Chester et al., 2018; Chlebowski et al., 2017; Haberer and Silversides, 2019; Mosca et al., 2007; Whayne and Mukherjee, 2015). Furthermore, current studies suggest that a sex-specific program controlled genetically through the X or Y chromosome can function outside of the sex organs (Arnold, 2017, 2019; Naqvi et al., 2019; San Roman and Page, 2019; Snell and Turner, 2018). Though it is clear that hormones play a critical role in cardiac disease, the mechanisms underlying sex differences in cardiac homeostasis and disease remain unexplained.

To define the mechanisms of cardiac sex differences, we initiated a directed, quantitative, proteomics-based approach to identify proteins, protein complexes, and protein pathways differentially expressed in adult and embryonic, male versus female, mouse heart tissue (Dorr and Conlon, 2019; Slagle and Conlon, 2016). Our findings showed that cardiac sex disparities occur through a post-transcriptional mechanism that establish sex differences through a set of proteins and protein pathways that act by hormonal and non-hormonal genetic mechanisms.

We have leveraged the power of the Collaborative Cross (CC) as a surrogate for human diversity, identifying the proteins, protein complexes and protein pathways that are conserved and those that diverge between males and females across heterogeneous populations. We report that the onset of sex-biased protein expression, is detected in cardiomyocytes at the earliest stages of heart development; a period shortly after their fate is determined and preceding mammalian gonadal development. Taken together, these studies elucidate mechanisms that initiate and establish cardiac sex disparities.

## RESULTS

### Cardiac sex differences are established by post-transcriptional mechanisms

To investigate the molecular basis of cardiac sex differences, we used an unbiased systems-based approach to characterize differential male and female cardiac transcripts in adult C57BL/6J mouse hearts (RNA-seq) (Fig. 1A). Analyses identified 16,160 genes, of which 223 showed differential sex expression, with 105 upregulated in male hearts and 118 upregulated in female hearts (Fig. 1B and Table S1). Pathway analysis revealed an enrichment of processes associated with male immune responses, while females showed enrichment in cilium, septation and trabeculation processes (Fig. 1C). These data suggest that fundamental differences in gene expression exist in male and female heart tissue.

**Figure 1.**
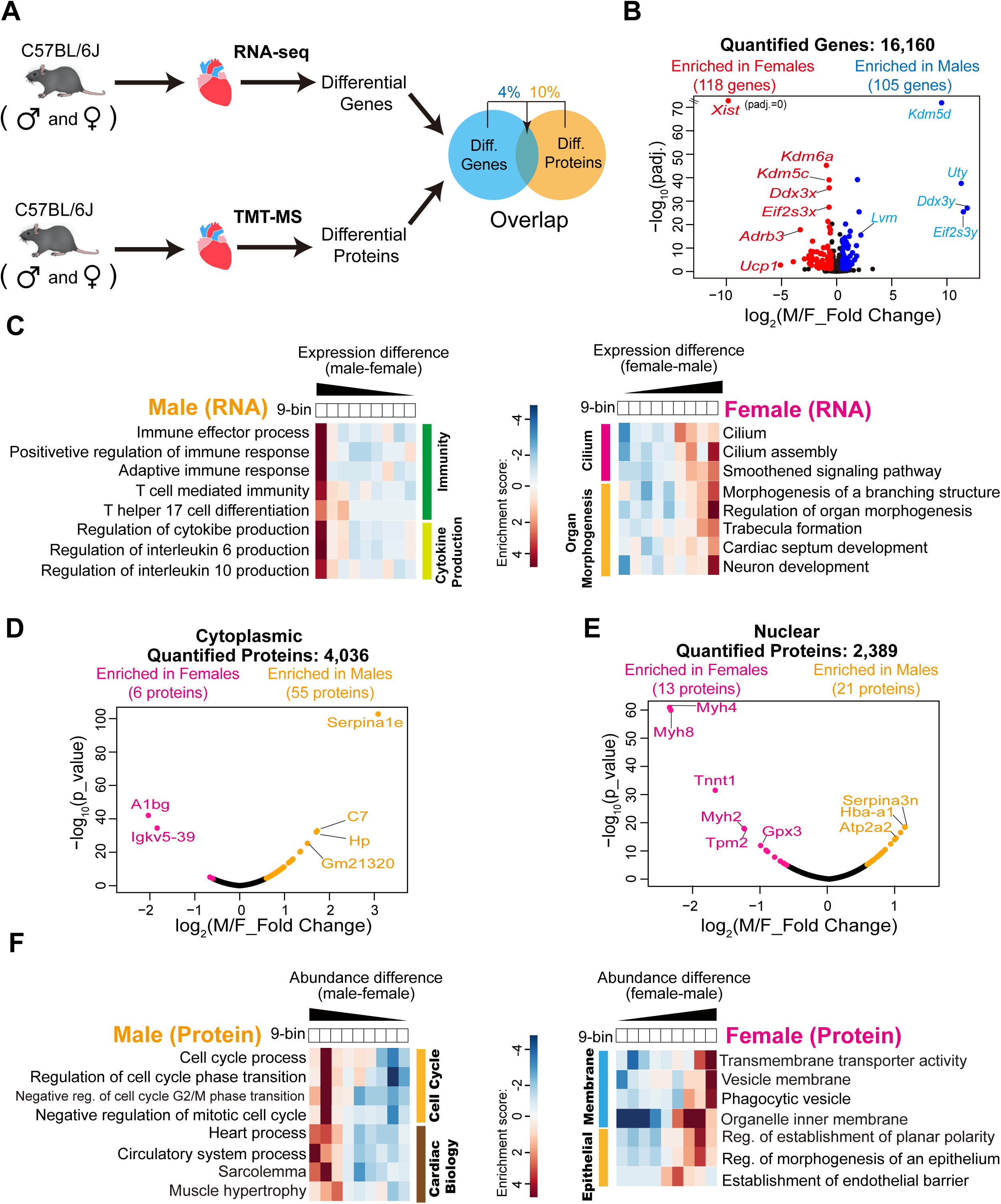
Cardiac transcriptome and proteome in C57BL/6J mouse. (A) Experimental overview of the transcriptomics and proteomics analyses in C57BL/6J (C57) mouse hearts from both sexes. (B) Volcano plot of RNA-seq in C57 mouse hearts showing log_2_ (Male/Female_Fold Change) plotted against the –log_10_ (padj.) (padj.: adjusted p_value) as determined by DESeq2. Genes with log_2_ (Male/Female_Fold Change) > 0.5, padj. < 0.05 are enriched in male hearts, blue dots; log_2_ (Male/Female_Fold Change) < -0.5, padj. < 0.05 are enriched in female hearts, red dots. Representative genes in each sex are labeled in the plot. N=4 per sex. (C) Representative enriched gene ontology (GO) terms (biological processes) of identified genes from C57 mouse hearts. Terms are divided into male-enriched (left) and female-enriched (right). Color bars: enrichment score. (D, E) Volcano plots of TMT-MS in C57 mouse hearts in each cellular fraction (d: cytoplasmic, e: nuclear) showing log_2_ (Male/Female_Fold Change) plotted against the –log_10_ (p_value). Proteins with log_2_ (Male/Female_Fold Change) > 0.59, p_value < 0.01 are enriched in male hearts; log_2_ (Male/Female_Fold Change) < -0.59, p_value < 0.01 are enriched in female hearts. N=2 per sex. (F) Representative enriched GO terms (biological processes) of identified proteins from C57 mouse hearts. Color bars: enrichment score. See also Figure S1 and Tables S1 and S2.

Studies have demonstrated that RNA processing plays an essential role in determining protein expression levels (Jordan et al., 2019). To determine if male-female cardiac mRNA differences are related to changes in protein abundances, we conducted quantitative mass spectrometry (MS) analyses (Fig. 1A). To increase depth, we separated heart tissue into cytoplasmic and nuclear fractions and then performed multiplexed tandem mass tagging mass spectrometry (TMT-MS). From the results, we quantified a total of 4,797 cardiac proteins. Pairwise comparisons indicated that 55 cytoplasmic proteins were enriched in males, while only 6 cytoplasmic proteins were higher in females (Fig. 1D and Table S2).

Twenty-one and 13 nucleic proteins had higher abundances in males and females, respectively (Fig. 1E and Table S2). Strikingly, none of the cardiac proteins displaying differential levels based on sex correspond to any of the differentially expressed mRNAs examined. Moreover, cardiac proteins displaying sex disparity were associated with disparate biological processes versus those at the RNA level (Fig. 1, C and F), with males showing enrichment in cell-cycle and heart processes (Fig. 1F, left) and females showing enrichment in membrane and epithelial-related pathways (Fig. 1F, right). These results imply that sex disparities in cardiac protein expression occur post-transcription. To test this hypothesis, we compared the male and female proteomes to our corresponding RNA-seq data. Only 4% of the RNAs that displayed 1.44-fold changes exhibited corresponding changes at the protein level (Fig. 1, B, D, and E and Fig. S1). These findings strongly suggest that sex differences in cardiac protein expression are established through post-transcriptional mechanisms.

### Conserved cardiac sex protein, protein complexes, and biological processes in heterogeneous populations

In addition to sex biases, cardiac disease states vary in frequency by ethnicity (van der Linde et al., 2011). To identify the proteins and protein pathways for which cardiac sex differences are conserved across heterogenous populations, we conducted TMT-MS analyses in male and female hearts across all 8 founding mouse strains of the Collaborative Cross (CC) (Fig. 2A and Fig. S2) (Churchill et al., 2004; Threadgill et al., 2011). The CC is a panel of recombinant inbred mouse strains derived from 8 founder strains that has greater diversity than the total human population and thus, serves as a surrogate for human variation (Churchill et al., 2004; Threadgill and Churchill, 2012; Threadgill et al., 2011). In total, we quantified 4,797 unique proteins across all 8 founding strains (Fig. S3 and Tables S3 and S4). Consistent with reported variabilities in gene expression between mouse strains (Fontaine and Davis, 2016; Hunter, 2012), we observed diversity in sex-biased proteins and protein pathways by strain (Figs. S4 and S5). For example, mouse strains PWK/PhJ and CAST/EiJ exhibit male bias in cardiac proteins expressed at the G2M checkpoint, while NZO/ HlLtJ and NOD/ ShiLtJ strains show female bias (Fig. S5A). A second example, strain 129S1/SvImJ shows male bias for oxidative phosphorylation, while all others exhibit female bias (Fig. S5B). Together, these data demonstrate significant differences between mouse strains in the cardiac proteins and pathways where we previously showed sex-biased expression. Given the well-established recognition of sex differences in cardiac surgical models (Doetschman, 2009; Fontaine and Davis, 2016; Hunter, 2012), our findings should provide insight regarding the best strain(s) for further investigation.

**Figure 2.**
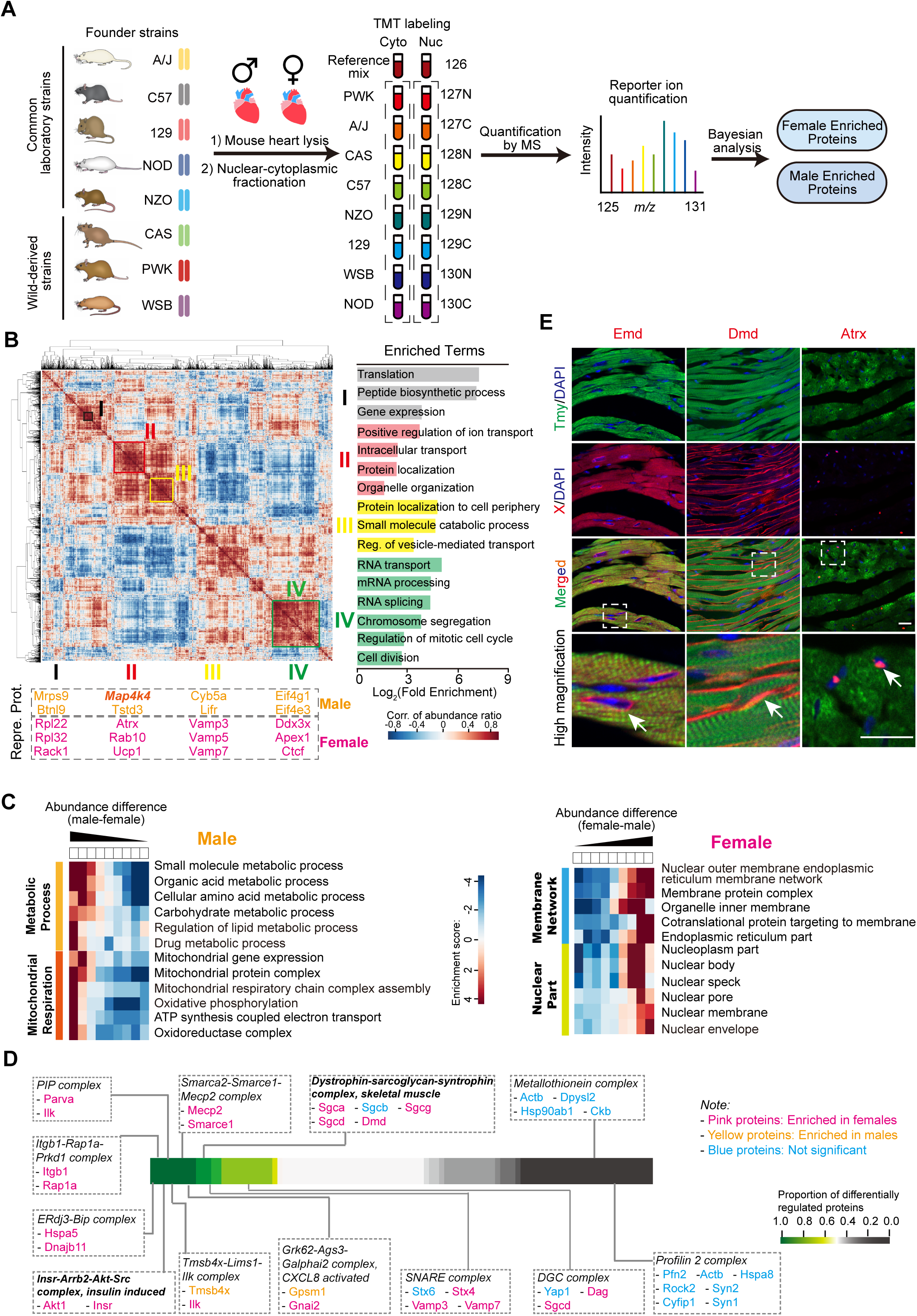
Global cardiac proteome in the founder strains of the Collaborative Cross mice. (A) Experimental design and TMT-MS proteomic workflow for the hearts of 8 founder strains. N=2 per strain per sex. (B) Correlation matrix of sex differences across all eight founder strains between proteins. Protein abundance ratios (log_2_ (Male/Female)) in eight strains were concatenated for each protein, and the correlation of the ratios were calculated between proteins. Color bar: correlation of abundance ratio (log_2_ (Male/Female)). 4 representative clusters are labeled in the heatmap (I - IV), representative enriched GO terms (FDR < 0.05, Fold Enrichment > 2, protein numbers involved in each term are more than 10) and representative proteins (Repre. Prot.) in each cluster are shown, respectively. (C) Representative enriched GO terms (biological processes) of identified proteins from the 8 founder strains hearts of both genders. Color bar: enrichment score. (D) Differential protein complex analysis. Representative identified protein complexes from the 8 founder strains hearts are shown in dashed line squares. Proteins in pink: enriched in females; in yellow: enriched in males; in blue: not significant. Color bar: proportion of differentially regulated proteins. (E) Representative immunohistochemistry (IHC) fluorescence staining images in adult female C57 heart sections. Cardiomyocytes are stained by Tropomyosin (green), protein of interests are stained with Emd (Emerin), Dmd (Dystrophin), or Atrx antibody (red, white arrows), nucleus are stained by DAPI (blue), scale bar: 25 μm. High magnification images showing areas in white dashed squares. Scale bars: 50 μm. See also Figures S2-S6 and Tables S3-S6.

In addition to identifying diverging proteins and pathways by strain in the CC samples, we used Bayesian inference to gain a broader understanding of the proteins that are conserved across strains in their sex-biased protein expression (Sjolander and Vansteelandt, 2019). This conditional probability approach models estimates of the contribution of a single protein across a population and, thus, allows us to detect small differences that are statistically significant; i.e., this approach classifies the proteins that are conserved in male-female abundance across the CC strains but not necessarily in any one strain itself. Using this approach, we identified a group of 1,379 differential male-female cardiac proteins across CC strains: 1,108 cytoplasmic and 271 nuclear proteins (Fig. S6). Proteins were annotated into pathways associated with peptide synthesis and gene expression (cluster I), intracellular transport and protein localization (cluster II), protein localization to cell periphery (cluster III), and RNA processing (cluster IV) (Fig. 2B and Table S5). In general, the conserved cardiac pathways found to exhibit greater enhanced bias in males are involved in metabolic processes (Fig. 2C, left), whereas conserved pathways exhibiting female-bias are associated with the architectures of the nucleus and endoplasmic reticula (Fig. 2C, right). These groups of conserved, differentially regulated proteins include proteins previously reported to be associated with cardiac sex-biased disease states (Mosca et al., 2011a; Peters et al., 2019). For example, decreases in Map4k4 protein levels have been associated with insulin-resistance, obesity, and atherosclerosis, all prevalent and often lethal female-biased disease states (Fiedler et al., 2020). Our results show consistently higher Map4k4 expression levels in male hearts (Fig. 2B). Taken together, these findings identify a conserved set of proteins and protein pathways across CC strains, a subset of which appear to contribute to male-female disease bias.

Proteins assemble and function in multi-component complexes. Consistent with female predisposition for upregulation in protein expression levels for cytoplasmic proteins (772 vs 336) and nuclear proteins (218 vs 53) (Figs. S6, C and D and Table S6), we found significant enrichments in protein complexes that are largely female-biased. For example, we observed female upregulation in proteins comprising the Dystrophin complex (Fig. 2D), a complex that, when mutated, leads to sex differences in the pathways of muscle wasting (Arnold et al., 2001; Giannakoulas et al., 2017; Maric-Bilkan et al., 2016; Rosa-Caldwell and Greene, 2019). A second example is the insulin induced pathway protein complex which was also upregulated in females; this pathway is associated with female biased obesity and cardiac disease (Arnold et al., 2001; Giannakoulas et al., 2017; Investigators et al., 2012; Maric-Bilkan et al., 2016; Peters et al., 2019). Collectively, our studies utilizing CC strains have identified and characterized a set of proteins, protein complexes and protein pathways that are conserved across a heterogeneous population.

### Cardiomyocytes exhibit sex-biased disparities in protein expression

Using our adult whole-heart samples, we sought to determine which of the heart’s multiple cell types express the sex-based differences in protein abundance that we identified. Western blot analyses confirmed that Atrx, Ddx3x, and A1bg protein levels are upregulated in females (Fig. S6F); we further found Atrx with Emd and Dmd to be co-expressed with Tropomyosin in the cardiomyocyte compartment of adult hearts (Fig. 2E). Therefore, our data demonstrate that cardiomyocytes can differentially express male-female proteins.

### Cardiac sex-bias protein expression is established by hormonal and non-hormonal mechanisms

We next sought to identify the mechanisms leading to sex-biased protein expression. Studies have postulated that sex differences can be established by hormonal mechanisms. However, recent work has also implied that cardiac sex differences may be regulated by inequalities in sex chromosome compliment (Arnold et al., 2017; Naqvi et al., 2019; San Roman and Page, 2019; Snell and Turner, 2018). To test the hypothesis that sex disparities in cardiac protein expression levels are due in part to sex chromosomal mechanisms, we quantified protein levels in adult cardiac tissue derived from the Four Core Genotypes (FCG) mouse model. The FCG model takes advantage of the deletion of the testis-determining *Sry* gene from the Y chromosome, and the insertion of *Sry* onto an autosome (Burgoyne and Arnold, 2016; Snell and Turner, 2018). Testes develop in mice with *Sry*, but gonadal sex is uncoupled from sex chromosome complement (XX vs. XY) (Fig. S7). Using the FCG model in 3 independent matings in the C57BL/6J background, we derived mice of all 4 genotypes and conducted TMT-MS analyses of the respective heart tissues (Fig. 3A). We measured 3,871 proteins in each of the 4 FCG genotypes. K-means clustering analysis identified nine clusters of proteins that showed significant differences between at least two of the 4 genotypes (Fig. 3B and Table S7). Our results indicate that 519 proteins segregate with ovaries and testes and thus are hormonally controlled; they segregate depending on the presence/absence of *Sry* (i. e. *Sry*-dependent) (Fig. 3C), while 159 proteins co-segregate with sex chromosomes (XX versus XY) (Fig. 3D). Taken together, our results strongly imply that sex differences in cardiac tissue occur via both hormonal and sex chromosome complement mechanisms.

**Figure 3.**
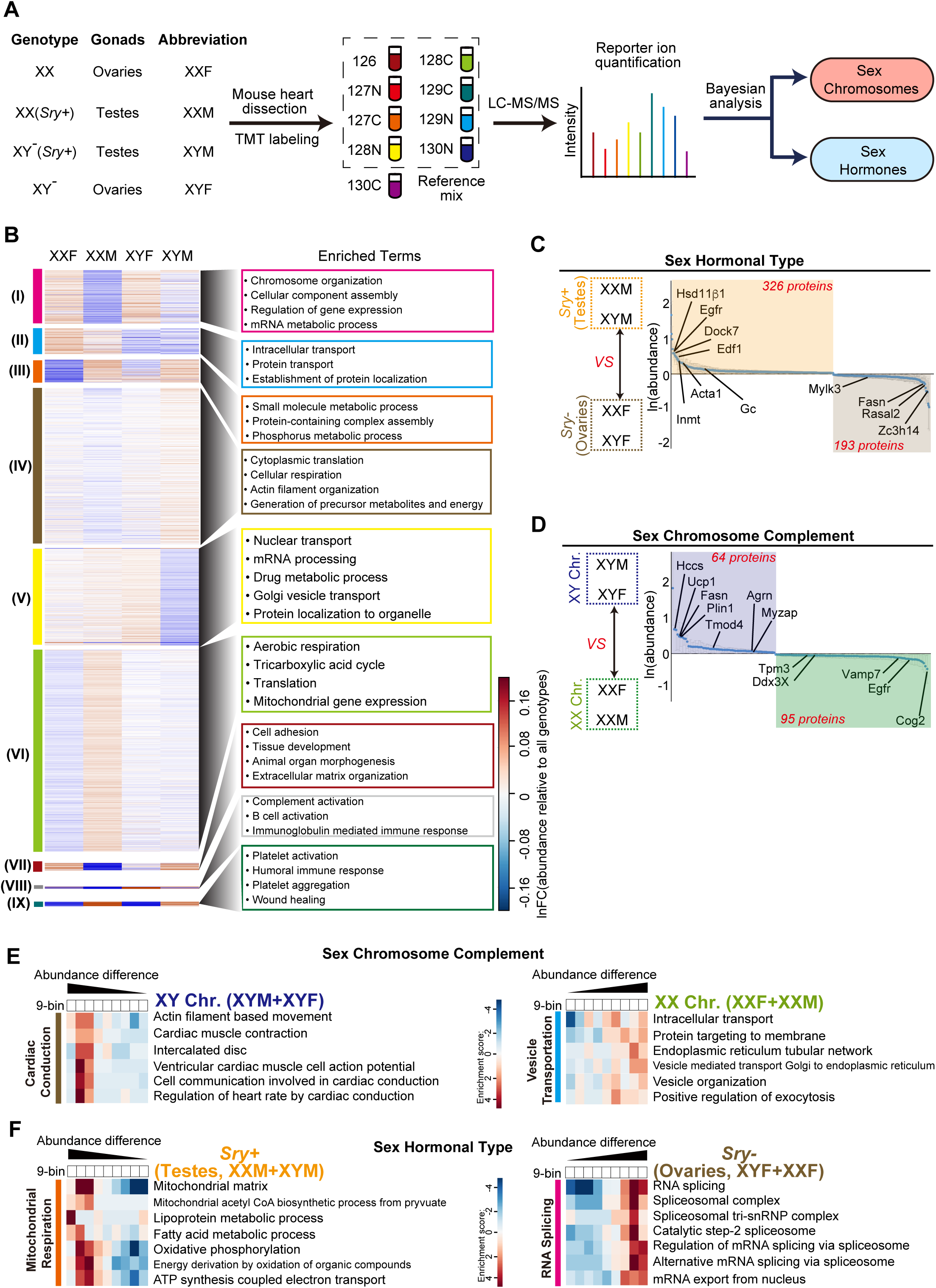
Cardiac proteome in the Four Core Genotype (FCG) mouse model. (A) Experimental design and TMT-MS proteomic workflow for the FCG mouse hearts. N=3 per genotype. Abbreviations (XXF, XXM, XYM, and XYF) are used to denote each genotype hereafter. Y^-^: Y chromosome deleted for *Sry. Sry*+: *Sry* transgene inserted into chromosome 3. (B) K-means clustering displayed as a heat map of the identified proteins from the FCG mouse hearts. Color bar: ln (abundance relative to all genotypes). Representative enriched GO terms (FDR < 0.05, Fold Enrichment > 2. For clusters I – VII, protein numbers involved in each term are more than 10; for clusters VIII and IX, protein numbers involved in each term are more 3) in each cluster are shown in specific frames. (C, D) Waterfall plots of differential protein abundance driven by sex hormonal types (*Sry*+/Testes (XXM and XYM) versus *Sry*-/Ovaries (XXF and XYF)) (C) and by sex chromosome complement (XX (XXF and XXM) versus XY (XYF and XYM)) (D) based on Bayesian inference analyses. Displayed proteins are those that whose 2.5% and 97.5% quantiles of posterior distribution (central 95% Bayesian credible interval) are both greater or less than 0. The mean of the posterior distribution of each protein and its credible intervals are plotted. Proteins with ln (XY/XX (or *Sry*+/*Sry*-)_abundance) > 0 are driven by XY chromosomes (XYM and XYF) or *Sry*+ (XXM and XYM), otherwise driven by XX chromosomes (XXF and XXM) or *Sry*- (XXF and XYF). Protein numbers are shown in each effect, and proteins of interest are labeled. (E, F) Representative enriched GO terms (biological processes) for sex chromosomes (E) and sex hormones (F) controlled proteins from the FCG mouse hearts. Color bars: enrichment score. See also Figures S7, S8 and Tables S7, S8.

Consistent with previous studies, we identified a significant number of proteins under hormone control (i.e., Egfr, Hsd11, Azgp1, Pin4) (Fig. 3C, Table S8). Of the 159 proteins controlled by sex chromosome complement, 64 are driven by XY chromosomes (i.e., Hccs, Ucp1, Ca3, Fasn), and 95 are driven by XX chromosomes (i.e., Vamp7, Cog2) (Fig. 3D and Table S8). Given observed differences in electrical conduction in male and female hearts (Arnold et al., 2001; Macfarlane, 2018) and the observation that men have higher rates of atrial fibrillation (Linde et al., 2018), it is interesting that the most prominent category in the XX-XY chromosome complement-dependent pathway is composed of major determinants in cardiac conduction processes (Fig. 3E, left). This implies that a non-hormonal pathway controls sex differences in cardiac conduction. Overall, these findings demonstrate sex differences in cardiac tissue are controlled by both hormonal and sex chromosomal mechanisms (Figs. 3, E and F)

### Cardiac sex differences are established prior to gonad formation

Congenital heart disease (CHD) is the most common of the congenital malformations. A significant portion of CHD states are developmental in origin and sex biased. To determine when, during embryogenesis, heart tissue displays sex disparities in protein expression, we utilized mice because their gonads do not differentiate as testes or ovaries until late in the second trimester (E11) (Garcia-Moreno et al., 2018; Nef et al., 2019), though mice have a well formed and beating heart prior to that time (Bruneau, 2003). We set up timed mating of C57BL/6J mice, dissected hearts at E9.5 from individual embryos, and then sexed the carcasses. Male and female hearts were pooled (N=3 for each sex, 20 hearts per sex) and we conducted label-free quantitative proteomics on each (Fig. 4A). A total of 7,433 proteins were quantified (Table S9), of which 335 and 334 displayed relatively higher levels in male E9.5 hearts and female E9.5 hearts, respectively (Fig. 4B and Table S10).

**Figure 4.**
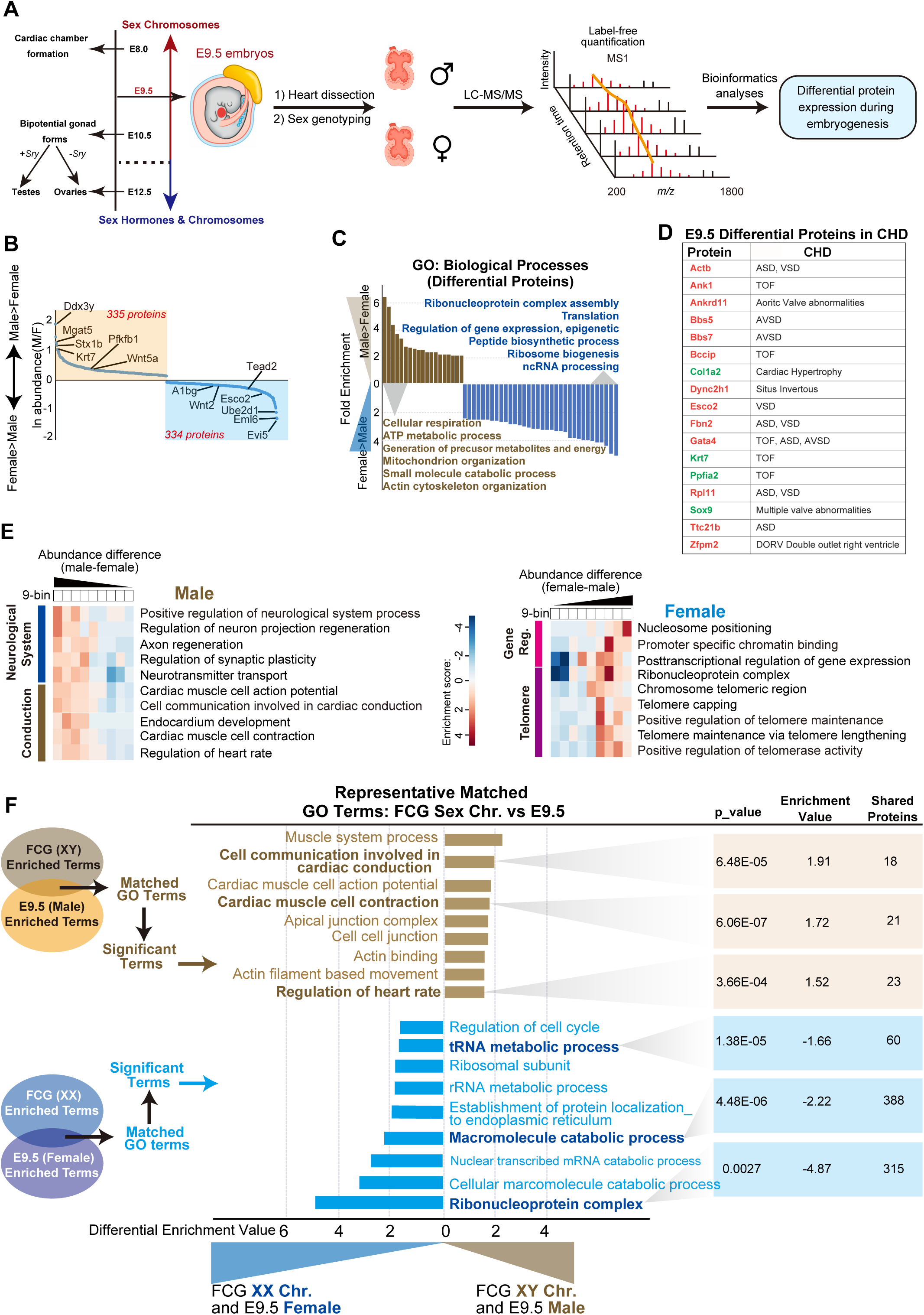
Cardiac proteome before sex gonads formation. (A) Experimental design and LC-MS/MS proteomic workflow for the E9.5 embryonic hearts. Three biological replicates per sex, 10 hearts per sex per replicate. (B) Waterfall plot of differential protein abundance between male and female in E9.5 embryos. (C) Representative enriched GO terms (FDR < 0.05, Fold Enrichment > 1.9, protein numbers involved in each term are more than 10) for the differential proteins in (B). The top 6 enriched terms in each sex are presented. (D) 17 differential proteins between males and females at E9.5 were associated with cardiac disease states that display sex disparity. Green proteins are enriched in males, red proteins are enriched in females. ASD: Atrial Septal Defects; VSD: Ventricular Septal Defects; TOF: Tetralogy of Fallot; AVSD: Atrioventricular Septal Defects. (E) Representative enriched GO terms of identified proteins from the E9.5 embryonic hearts. Color bar: enrichment score. (F) Representative matched GO terms between FCG_sex chromosome effect and E9.5 embryonic hearts datasets. Top 200 enriched GO terms in each dataset are used to perform the matching (FCG_XY vs E9.5_Male, FCG_XX vs E9.5_Female). Matched terms are further filtered by the following cutoffs: p_value < 0.05 and abs(enrichment value) > 1.5 in E9.5 datasets, and these selected GO terms are defined as the Significant Terms, suggesting that they are conservatively controlled by XX or XY chromosomes in embryonic and adult stages. Representative Significant Terms of “FCG_XY vs E9.5_Male” are shown in brown, and “FCG_XX vs E9.5_Female” are shown in blue. See also Tables S9, S10.

To determine if sex-biased protein expression is temporally regulated or if sex differences in expression persist into adulthood, we compared our results at E9.5 to those obtained in an adult. We found that a small subset of protein and protein processes were temporally conserved, including enhancement in males for proteins associated with conduction and contraction and enhancement in females for proteins associated with metabolic processes (Fig. 4F). However, most of the cardiac sex-differentiated proteins at E9.5 displayed temporal differences by adulthood (Fig. 4, B-E). Given that our analyses were conducted prior to gonadal development and subsequent sex hormone expression, our findings strongly imply that male-female cardiac protein differences are controlled by sex chromosomes at this stage. Furthermore, given that *Sry* is not expressed in the embryo prior to or at this stage, our findings suggest that sex differences at E9.5 are regulated by *Sry*-independent pathways that function before gonadal development.

## DISCUSSION

We performed a systems-based approach to identify quantitative molecular differences between male and female adult hearts at both RNA and protein levels. Contrary to current thinking, we show that cardiac sex differences are established by post-transcriptional mechanisms. We further find that heart tissue displays diffences in protein expression prior to expression of the testes determining factor *Sry*. These finding may explain the observation that a number of CHD disease states are sex biased even if the causative gene does not dispaly sex biased experession, e.g. GATA4.

Our studies further found that only 3% of all cardiac proteins showing significant differences in expression between males and females mapped to genes on the X-chromosome, and none mapped to the Y chromosome. From these results, we anticipated that the initiating events are regulated transcriptionally and then propagated and maintained by post-transcriptional RNA mechanisms, (i.e., alternative polyadenylation and/or transcription start site, etc.). These events, in turn, result in male-female alternative translation efficiency, stability, and/or processing that lead to sex-differentiated protein expression in cardiac tissues.

### Cardiac sex disparities occur through hormonal and non-hormonal genetic mechanisms

Clinical investigations of the mechanisms behind sexual dimorphism have challenged the paradigm that sex hormones account for all differences in male versus female prevalence, treatment, and survival of human disease. Consistent with these findings, we have used our approaches to uncover and dissect the distinct hormonal and genetic pathways associated with sex differences in cardiac protein expression. We demonstrate that, contrary to present models, male-female cardiac differences are not solely controlled by hormones but are also controlled by non-hormonal genetic pathways. We further find evidence that some cardiac processes involve a combination of sex chromosome and gonad regulation, including processes associated with the extracellular matrix, cardiac action potential, and mitochondrial function (Fig. S8). It is possible that these pathways are regulated by both sex-chromosomal and gonadal mechanisms. Alternatively, these pathways may be driven by sex chromosome regulation but counteracted by opposing sex hormones, thus canceling the effect of each (i.e., XYM would be equal to XXF). In the later scenario, we predict the patient phenotype may be similar between males and females for disease states involving these proteins: however, the patient phenotype may arise via different mechanisms, such that treatment of the disease may vary between males and females.

### Sex differences in the cardiomyocyte lineage are established prior to gonad formation

Congenital heart disease is the most prevalent and lethal form of birth defects in the US and Europe (Dolk et al., 2010; Heron et al., 2009). A significant portion of these disease states is sex biased (Haberer and Silversides, 2019; Kararigas et al., 2014; Prabhavathi et al., 2014). Our findings demonstrate that the onset of sex-biased protein expression in cardiomyocytes occurs at the earliest stages of heart development, shortly after cardiomyocyte fate is determined and prior to gonad formation. These findings are broadly consistent with studies that show sex-biased expression of transcription and epigenetic factors are maintained during differentiation of embryonic stem cells into cardiac precursors (Werner et al., 2017). Of the proteins we found differentially regulated among males and females at E9.5, 17 were associated with congenital cardiac disease states that display sex disparity (Fig. 4D).

However, we have found no reports of these proteins displaying sex differences in expression during development. Five of these proteins are encoded by genes known to cause Tetralogy of Fallot (TOF), a severe congenital heart defect that, even when taking into account risk factors, preferentially effects males: Krt7, Bccip, Ank1, Ppfia2, and Gata4 (Jin et al., 2017; Lahm et al., 2015). The finding that Gata4, an extensively studied cardiac transcription factor, is up-regulated in female E9.5 hearts is of additional clinical importance, given that mutations in Gata4 are also causally associated with two other prevalent sex-biased disease states: atrial septal defects and atrioventricular septal defects (Zhou et al., 2017).

Together with our results from the FCG model, our findings from E9.5 male-female heart tissue imply that X-linked genes acting via a dosage-specific mechanism initiate cardiac sex differences. This hypothesis is supported by the observation that over 70% of Turner Patients (female, XO) have congenital heart disease (Granger et al., 2016; Koenraadt et al., 2019; Mortensen et al., 2010). Testing this hypothesis and identifying the genes on the X-chromosome that initiate sex disparities in a cardiac specific manner will clarify our understanding of how X-linked genes function in a tissue-specific manner. We further hypothesize that these mechanisms not only regulate cardiac male-female differences but also apply to sex disparities observed in many other disease states, including cancer, dementia, chronic kidney disease, obesity, and autoimmune disease (Babapour Mofrad and van der Flier, 2019; Channappanavar et al., 2017; Christou et al., 2019; Kadioglu et al., 2011; Leung, 2020; Liang et al., 2017; Luders et al., 2009; Snell and Turner, 2018; Zhu et al., 2000).

## METHOD DETAILS

### Mouse

The founders of the Collaborative Cross (CC) were obtained from JAX, and housed at the Division of Laboratory Animal Medicine, University of North Carolina at Chapel Hill (UNC-CH). The FCG mice were maintained by Arthur Arnold’s lab at University of California, Los Angeles. Mice were sacrificed at 8 weeks, and hearts were perfused with 1×PBS and dissected, snap frozen for proteomics or western blot analysis, or perfused with 4% paraformaldehyde (PFA)/0.1% Tween-20/PBS for immunohistochemistry staining, or dissected and immediately homogenized in Trizol for RNA extraction for RNA-seq analysis. Research was approved by the Institutional Animal Care and Use Committee at UNC-CH and conformed to the Guide for the Care and Use of Laboratory Animals.

### RNA-sequencing analysis

RNA extraction from individual heart (n=4 per sex) was performed as previously described (Wilczewski et al., 2018), Poly-A selected RNA-seq libraries preparation and sequencing reactions were conducted at GENEWIZ, LLC. (South Plainfield, NJ, USA). Samples were run on a HiSeq2500 (Illumina) with 2×150 bp paired end reads.

Sequence reads were trimmed to remove possible adapter sequences and nucleotides with poor quality using Trimmomatic v.0.36. The trimmed reads were mapped to the Mus musculus GRCm38 reference genome available on ENSEMBL using the STAR aligner v.2.5.2b. Unique gene hit counts were calculated by using featureCounts from the Subread package v.1.5.2. The hit counts were summarized and reported using the gene_ID feature in the annotation file. Only unique reads that fell within exon regions were counted. After extraction of gene hit counts, the gene hit counts table was used for downstream differential expression analysis. Using DESeq2, a comparison of gene expression between the customer-defined groups of samples was performed. The Wald test was used to generate p-values and log_2_(fold changes). Genes with an adjusted p-value < 0.05 and absolute log_2_(fold changes) > 0.5 were called as differentially expressed genes for each comparison.

### Dissection and sex genotyping of E9.5 embryonic hearts

Each E9.5 heart was dissected in ice-cold PBS and snap frozen in liquid nitrogen individually, primers for genotyping the sex of each embryonic heart were listed in the KEY RESOURCES TABLE.

### Immunohistochemistry

Hearts from two male and two female C57BL/6J mice were fixed in 4% PFA/0.1% Tween-20/PBS at 4°C overnight, dehydrated by sucrose gradient and embedded in OCT followed by cryostat sectioning. Immunofluorescent staining was performed with antigen retrieval on 8 μm coronal cryosections as previously described (Dorr et al., 2015). Primary and secondary antibodies information were listed in the KEY RESOURCES TABLE. Immunohistochemistry images were captured on a Zeiss LSM 700 laser scanning confocal microscope. ImageJ (NIH) was used for image analysis and standard image processing.

### Western blot

Snap frozen hearts (3 males and 3 females) were homogenized using a mortar and pestle in liquid nitrogen. Lysed the heart homogenate on ice for 30 min in RIPA buffer (50 mM Tris8.0, 0.5% Sodium Deoxycholate, 150 mM NaCl, 1% Triton-X 100, 0.1% SDS) with protease inhibitors. Quantification of protein concentration in total lysates was performed by using BCA Protein Assay Kit (Thermo #23225). Equal protein amount (40 μg) from each sample was loaded for SDS-PAGE gel electrophoresis. Western blots were blocked with 5% milk/TBST at room temperature for 1 hr, and then probed with specific primary antibodies overnight at 4 °C and followed by probing with secondary antibodies at room temperature for 1 hr.

Antibody-antigen complexes were visualized using an ECL Western Blotting Analysis System (Amersham). Primary and secondary antibodies information were listed in the KEY RESOURCES TABLE. Quantifications of protein bands from western blot films was performed with ImageJ, relative expression of each target protein were determined by the ratio to GAPDH.

### Mass spec sample preparation

32 founder strain hearts (8 founder strains, 2 male and 2 female replicates for each strain) and 12 FCG hearts (4 FCG genotypes, 3 replicates for each genotype) were prepared for mass spectrometry analysis with TMT 10-plex labeling. E9.5 hearts (3 male and 3 female replicates, 20 pooled hearts for each) were prepared for label-free quantitation.

Subcellular fractionation was performed on the founders hearts with 2 biological replicates per strain. Briefly, hearts were diced, snap frozen in liquid nitrogen, and then blended in a stainless-steel waring blender. Tissue powder was dissolved in a modified NIB-250 buffer, containing 10 mM Tris, pH 7.4, 250 mM sucrose, 1mM EDTA, 0.15% NP-40, 10 mM sodium butyrate, 1 mM PMSF, and 1X HALT protease and phosphatase inhibitor (Pierce). After gentle centrifugation at 400 rpm, the supernatant (cytosolic fraction) was transferred to a new low-bind tube and the pellet (nuclear fraction) was resuspended in lysis buffer (20 mM K-HEPES, pH 7.4, 110 mM KOAc, 2 mM MgCl2, 0.1% Tween-20, 1 mM ZnCl2, 1 mM CaCl2, 150 mM NaCl, 0.5% Triton x100, 1X HALT protease and phosphatase inhibitors). Following subcellular fractionation, lysates were reduced and alkylated with 25 mM tris(2-carboxyethyl)phosphine (TCEP, Pierce, Waltham, MA) and alkylated with 50 mM chloroacetamide (CAM, ThermoFisher Scientific, Waltham, MA) for 20 min at 70°C, methanol/chloroform precipitated, dried briefly in speed vac, and resuspended in 25 mM HEPES, pH 8.2. BCA assay was performed to estimate protein concentration, and 50 μg of protein was digested with sequencing grade trypsin (Promega) at 1:50 trypsin:sample protein amount for 14 hrs on a ThermoMixer at 37°C and 600 rpm. Samples were concentrated in a vacuum centrifuge to generate 50 mM HEPES, pH 8.2, adjusted to 20% ACN, and labeled with 10 μL of 15.4 μg/μL of the corresponding TMT 10-plex reagent for 1 hr at room temperature with shaking at 1000 rpm. The labeling reaction was quenched with 7 μL of 5% hydroxylamine, and the equal amount (2 μL) of labeled peptides were then mixed with 0.1% trifluoroacetic acid (TFA) (Pierce) to generate a test mix which was desalted by StageTip fractionation with Empore C18 filters (3M) (Rappsilber et al., 2007). Disks were first charged with a 100% acetonitrile (ACN) wash, followed by 2 conditioning washes with 0.1% TFA. Test mix samples were then applied to the StageTip, washed twice with 5% ACN with 0.1% TFA, and eluted with 70% ACN with 1% FA. Eluted samples were then dried in a vacuum centrifuge, resuspended in 5 μL of 2% ACN with 1% FA. Once the test mix was analyzed by mass spectrometry (described below), equal amounts of each TMT channel were mixed according to the derived ratios generated from the Proteome Discoverer 2.2 search of the data (described below).

The sex reversed FCG mouse hearts (3 biological replicates per genotype) were prepared in a similar manner as the founders samples, however, subcellular fractionation was not performed on these hearts. Isolated hearts were first minced into thin slivers and homogenized in 4 mL of 50 mM Tris-HCl, pH 8.0, 100 mM NaCl, 0.5 mM EDTA, and 1x HALT protease and phosphatase inhibitor cocktail (Pierce) with 40 strokes in a Tenbroeck homogenizer. Lysates were then transferred to 15 mL tubes and mixed 1:1 with the same buffer containing 4x SDS to generate a final SDS concentration of 2% SDS. These samples were then heated at 95°C for 5 min and then sonicated in a cup horn sonicator (1 sec pulses, medium power) individually for 20 seconds, repeated 3 times or until no insoluble material remained. Protein concentration was estimated using the BCA assay, 50 µg of protein for each sample was aliquoted and reduced and alkylated, methanol/chloroform precipitated, resuspended with 25 mM HEPES, pH 8.2, and digested with trypsin, as with the Founder line hearts above. TMT labeling and test mix desalting were performed as described earlier. The TMT mixes for the founders and FCG mice were subsequently desalted and fractionated with a Pierce High pH Reversed-Phase Peptide Fractionation Kit (Thermo #84868), as per the manufacturer’s protocol.

Embryonic day 9.5 mouse hearts were prepared for label free quantitation. Due to the low amount of protein extracted per embryonic heart (∼4-6 μg per heart), 20 mouse hearts were combined per replicate, with three biological replicates performed per sex. Embryonic hearts were lysed in 50 mM Tris-HCl, pH 8.0, 100 mM NaCl, 0.5 mM EDTA, and 5% SDS with sonication in a cup horn sonicator (1 sec pulses, medium power) individually for 20 seconds, followed by heating at 95°C for 5 min, repeated 3 times or until no insoluble material remained. Samples were then reduced and alkylated with 20 mM TCEP and 40 mM CAM for 20 min at 70°C. The SDS lysate was then acidified with phosphoric acid to generate a final concentration of 1.2% phosphoric acid. Samples were then digested with an S-Trap (ProtiFi) following the manufacturer’s protocol. Briefly, samples were adjusted to 90% methanol, applied to the S-Trap column, washed with S-Trap binding buffer (100 mM triethanolamine bicarbonate, pH 7.1, with 90% methanol), followed by the addition of 1 μg of trypsin in S-Trap digestion buffer (25 mM TEAB, pH 8), with digestion at 47°C for 1 hr.

Digested peptides were sequentially eluted from the S-Trap column with 40 μL of 25 mM TEAB, pH 8.0, 0.2% FA, then 50% ACN with 0.2% FA. These elution steps were combined, dried in a vacuum centrifuge, and then dried peptides were fractionated with Pierce High pH Reversed-Phase Peptide Fractionation Kit, as with the FCG samples.

### Basic reverse-phase fractionation

Each TMT mix for the founders was fractionated by StageTip fractionation with Empore C18 filters (3M) as previously described (Federspiel et al., 2019). Briefly, samples were dried in a vacuum centrifuge and resuspended in 1% TFA. 5 C18 filter disks were packed into a pipette tip, and were first charged with 100% ACN, equilibrated with 1% TFA, and then samples were applied to the disks. For test mix TMT runs, samples were desalted with 0.5% FA with 5% ACN in UHPLC water and eluted with 0.5% FA with 70% ACN in UHPLC water. For quantitative TMT samples, appropriate amounts of each TMT channel were mixed according to the derived quantitative values, and these mixtures were dried in a vacuum centrifuge, resuspended in 1% TFA, and then desalted and fractionated by StageTip fractionation. Disks were first charged with 100%, equilibrated with 1% TFA, samples were loaded onto disks, desalted with 5% ACN in UHPLC water, and samples were eluted in 20 fractions with sequential elution (20 μL each) ranging from 0.04 M Ammonium Hydroxide/8% ACN to 0.04 M Ammonium Hydroxide/46% ACN. The twenty collected fractions were concatenated to 10 fractions by combining fraction 1 and 11, 2 and 12, etc. For the FCG and E9.5 hearts, desalting and fractionation was performed with Pierce High pH Reversed-Phase Peptide Fractionation Kits (Thermo Scientific) to generate 8 fractions per sample, as per the manufacturer’s protocol. All fractions were dried to near dryness in a vacuum centrifuge and resuspended in 1% FA with 2% ACN in UHPLC water for liquid chromatography tandem mass spectrometry (LC-MS/MS) analysis.

## QUANTIFICATION AND STATISTICAL ANALYSIS

### Mass Spectrometry analyses

TMT labeled tryptic peptides (founder line strains and FCG strains) were separated with an EasyNano nLC1000 UHPLC (ThermoFisher Scientific) (running mobile phases: A (aqueous), 0.1% FA/H20; B (organic), 0.1% FA/97% ACN/2.9% H20) coupled online to an EASYSpray ion source and a Q Exactive HF (ThermoFisher Scientific). Samples were separated on ananocapillary reverse-phase PepMapC18 analytical column (75 µm by 500 mm; particle size 1.8 µm) (Thermo Scientific,) heated to 50°C with a 120-min linear gradient (6%-29% mobile phase B) at a flow rate of 250 nL/min. MS1 spectra were acquired over the m/z range from 400-1800 at a resolution of 120,000, an AGC of 3×106, and a Maximum Injection Time (MIT) of 30 ms. Data-dependent selection and fragmentation was performed on the top twenty most intense precursor ions by collision-induced dissociation and MS2 scans were acquired at a resolution of 45,000 with an AGC setting of 1×105, a MIT of 72 ms, and an isolation window of 0.8 m/z. Label-free analysis (E9.5 heart fractions) was performed in the same manner, except samples were separated with a 150-min gradient (5%-30% mobile phase B), and data-dependent acquisition was performed at a resolution of 15,000, with an AGC of 1×105, and a MIT of 150 ms.

### Database searching and reporter ion quantitation or label-free quantitation

For the founder and FCG strains, Proteome Discoverer 2.2.0.388 was used to search the LC-MS/MS data utilizing a fully tryptic Byonic search node in the processing workflow, searching against a combined Uniprot database containing M. musculus canonical protein sequences with common contaminants (17537 entries, downloaded 2018_08). A max of 2 missed cleavages were allowed, and peptides were required to have 5 ppm precursor accuracy and 10 ppm fragment accuracy. Peptides modifications included static cysteine carbamidomethylation, dynamic methionine oxidation, dynamic N-terminal methionine loss with N-terminal acetylation, dynamic asparagine deamidation, dynamic phosphorylation of serine, threonine, and tyrosine residues, and static TMT 10-plex labeling reagent on free peptide N-termini and on the □-nitrogen on lysine residues. The reporter ion quantifier node was utilized for TMT fragment ion quantitation. In the FCG samples, we observed a distinct batch effect by PCA, we observed a distinct batch effect by PCA, where one of the four litters occupied a distinct PCA space from the others. Thus, data acquired from channels 126, 128C, 129N, and 129C in the TMT plex were excluded.

For the E9.5 hearts, Proteome Discoverer 2.4.305 was used to search the LC-MS/MS data utilizing a fully tryptic SequestHT search node in the processing workflow, searching against a combined Uniprot database containing M. musculus canonical protein sequences with common contaminants (17537 entries, downloaded 2018_08). A max of 2 missed cleavages were allowed and peptides were required to have 5 ppm precursor accuracy and 0.02 Da fragment ion accuracy. Peptides modifications included static cysteine carbamidomethylation, dynamic methionine oxidation, dynamic N-terminal methionine loss with N-terminal acetylation, dynamic asparagine deamidation, and dynamic phosphorylation of serine, threonine, and tyrosine residues. Statistical analysis of male/female differential abundance was performed using the Background test in Proteome Discoverer.

### Gene Set Enrichment Analyses

Enrichment analyses were adapted from FIRE/iPAGE method (Goodarzi et al., 2009). The ratios of protein (RNA) abundance between males and females (M/F), or loge(protein abundance) derived from Bayesian analysis, were sorted from higher to lower. Sorted ratios were divided into N = 9 bins, which contain the same number of proteins (transcripts). C5 GO gene sets curated by MSigDB was used to annotate proteins (http://software.broadinstitute.org/gsea/msigdb/collections.jsp).

For each GO gene set, an array C of 2 × *N* was created, where N represents the number of bins. C(0,j) indicates the number of genes in the jth (*i* ≤ *j* ≤ *N*) bin but not included in the GO term being analyzed; C(1,j) represents the number of genes in the jth bin and included in the GO term. The mutual information (MI) of each bin in C was calculated using scikit-learn mutual_enrichment_score function, as 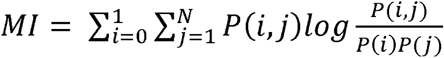, where 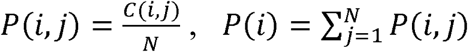, and 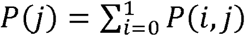. The MI scores represents the enrichment levels of the term being analyzed in each bin. To calculate z-scores for MI scores, we created a null distribution of the MI scores with 1,000 permutations with the same number of terms. Z-scores were defined as 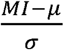, where *µ* and *σ* represent the mean and standard deviation, respectively.

To determine the level of enrichment, we defined the enrichment scores (*ESj*) of the jth bin as *s* × *log*_10_*p* − value of the hypergeometric distribution, where s represents 1 or -1 as described below. The p-value for the enrichment score was calculated by the hypergeom function of Scipy as *min* (*p*_*over*,_ *p*_*under*_), where 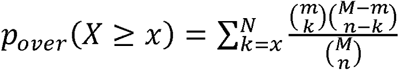, and 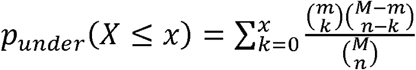. M, C, k and x represent 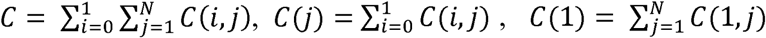, and *C* (1, *j*), respectively. If *p*_*over*_ > *p*_*under*_, s represents 1, otherwise -1. To estimate the degree of enrichment for male or female, we calculated the enrichment bias scores (*ES*_*bias*_), defined as 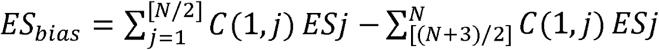. *ES*_*bias*_ were sorted from the highest to the lowest, and the top 200 terms were selected for z-score calculation and heatmap plotting. Heatmaps of the ES values were plotted with matplotlib and seaborn packages in Python.

### Bayesian inference analysis

Bayesian inference analyses were conducted using RStan in R (ver. 3.6.2). For the founders samples, we aim to obtain the posterior distributions of protein abundances contributed by sex and strain. We denoted the quantified protein abundance, average reference abundance, batch effect, sex effect, and strain effect as *µ*_*data*_, *µ*_*reference*_, *µ*_*batch*_, *µ*_*sex*_, and *µ*_*strain*_, respectively, and *µ*_*data*_ = *µ*_*reference*_ + *µ*_*batch*_ + *µ*_*sex*_ + *µ*_*strain*_. We used a normal distribution as a prior for batch effect, and uniform priors for reference average, sex effect, and strain effect. A flat uninformative prior was used for the standard deviation of data *σ*_*data*_ given no prior knowledge. A normal distribution was used to fit the posterior distribution: Norm *µ*_*data*_, *σ*_*data*_. No-U-Turn sampler (NUTS) was used for sampling, with 3 chains, 4000 iterations and 1000 warmup iterations. Proteins whose credible intervals are exclusively positive or negative are considered differential. For the FCG samples, sex and strain effects were replaced by chromosomal and gonadal effects, denoted as *µ*_*chrom*_ and *µ*_*gonad*_, and *µ*_*data*_ = *µ*_*reference*_ + *µ*_*batch*_ + *µ*_*chrom*_ + *µ*_*gonad*_. The same MCMC settings were used for this analysis as described above.

## Supporting information

Key Resource Table

## ACKNOWLEDGEMENTS

The authors would like to thank the Princeton University Proteomics Core Facility for mass spectrometry data analyses. This work was supported by grants R01HD089275 NIH /NHLBI and R01HL127640 NIH/NHLBI to F.L.C. and I.M.C.

## AUTHOR CONTRIBUTIONS

W.S., X.S., I.M.C., and F.L.C. designed and interpreted experiments. W.S., X.S., K.M.D., J.E.H., H.A.D., T.D.A., and X.C. performed the experiments. W.S., X.S., J.E.H., T.M.G., Y.H., J.D.F., Z.L.R., A.P.A., I.M.C., and F.L.C. analyzed data. W.S., X.S., J.E.H., T.M.G., A.P.A., I.M.C., and F.L.C. wrote the paper.

## SUPPLEMENTAL INFORMATION

### Supplemental Figure Legends

**Figure S1.**
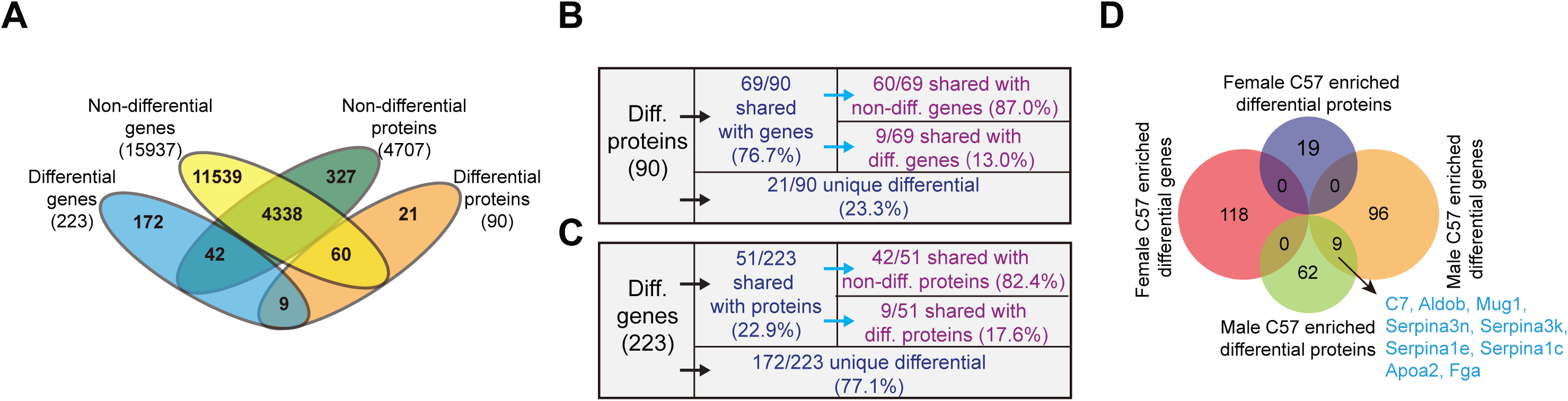
Overlap comparison of transcripts and proteins. (A-C) All identified transcripts and all characterized proteins are overlapped in terms of differential or non-differential. In the Venn diagrams (A), 223 differential genes are represented in blue, 90 differential proteins are represented in orange, 15,937 non-differential transcripts are represented in yellow, and 4,707 non-differential proteins are represented in green. (B, C) Breakdown tables of differential proteins (B) and genes (C). (D) Overlapping of differential genes and differential proteins in each sex. Overlapped proteins are indicated.

**Figure S2.**
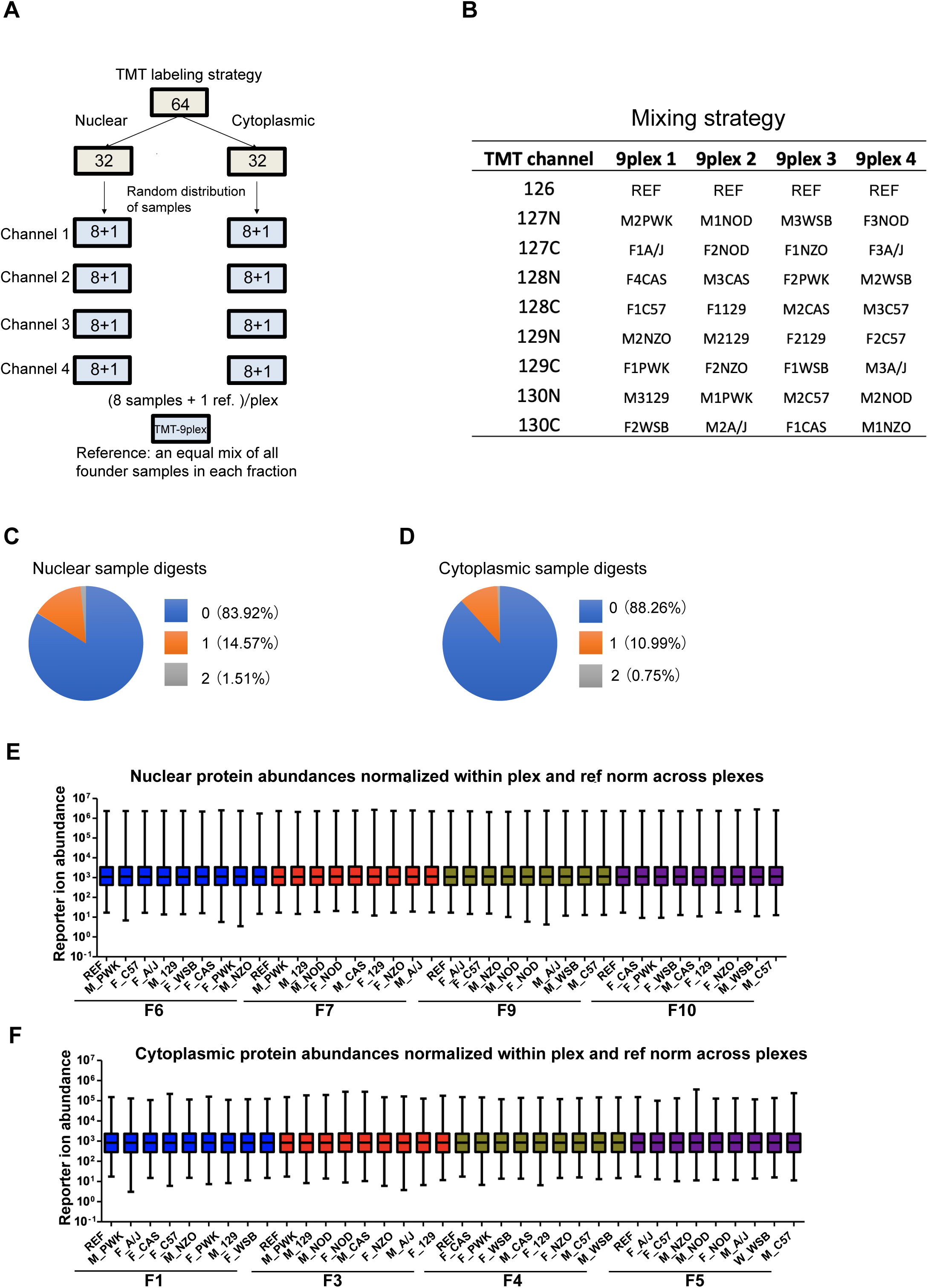
TMT labeling for quantification of mouse heart of founder strains. (A) Experimental design and TMT labeling strategy for nuclear and cytoplasmic samples. Nuclear and cytoplasmic fractions of mouse hearts were collected in duplicates (2 samples per sex per replicate) for all strains. The 32 samples were subdivided into 4 plexes with eight sample channels and 1 reference channel. (B) Mixing strategy for plex 1-4. Nuclear or cytoplasmic samples were randomly distributed into 4 plexes, and one reference channel (Channel 126) was used for subsequent normalization. (C, D) Statistics of trypsin miscleavage events in nuclear (C) and cytoplasmic (D) samples. (E, F) Protein normalization for nuclear (E) and cytoplasmic (F) proteins. Protein abundances were normalized within plex using median, and all protein abundances across plexes were normalized to the reference channels.

**Figure S3.**
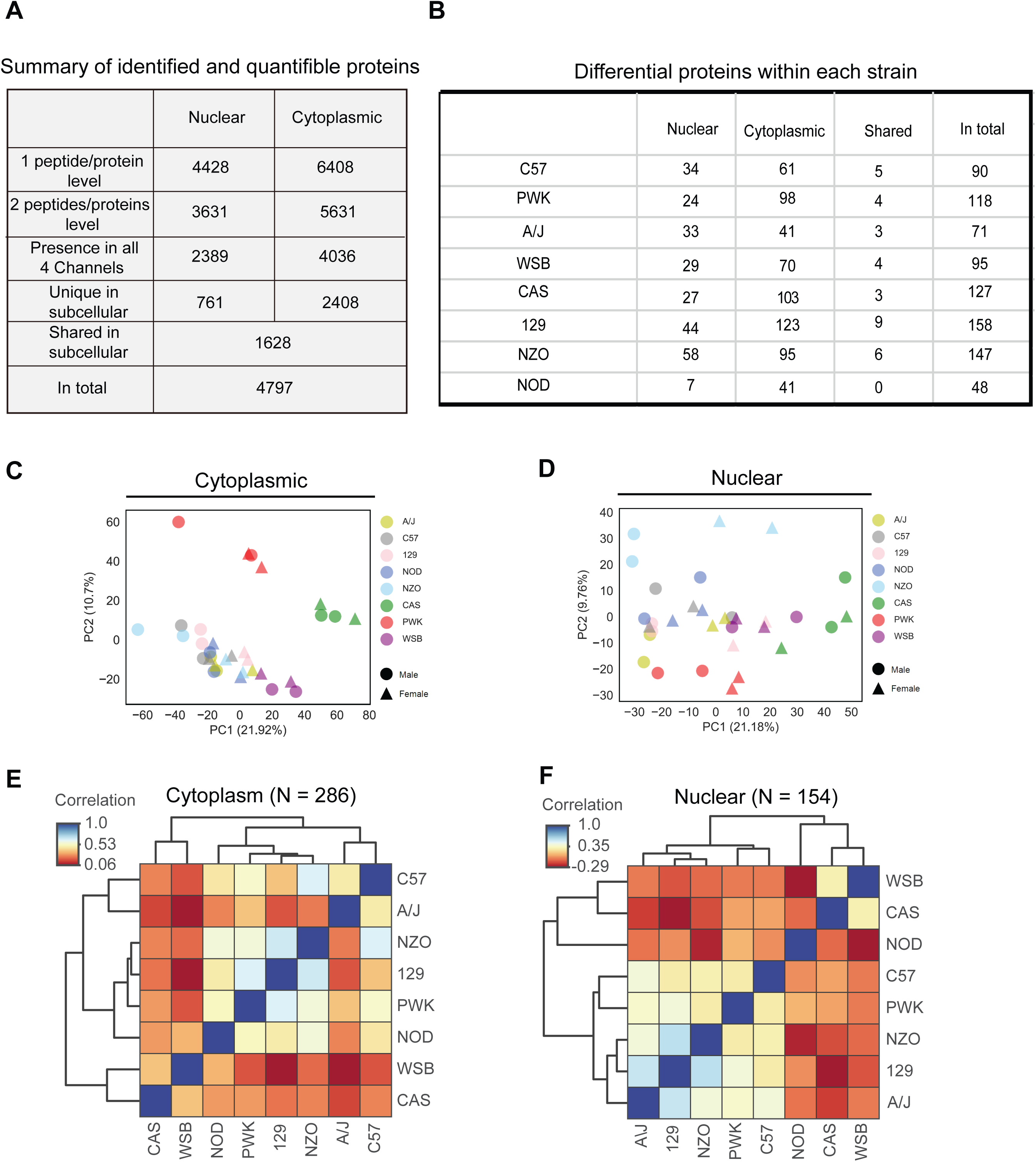
TMT-MS in founder strains. (A) Summary of quantified proteins in this assay. (B) Differential proteins based on pairwise comparisons in each strain. Proteins with log_2_ (Male/Female_Fold Change) > 0.59, p_value < 0.01 are enriched in male hearts; log_2_ (Male/Female_Fold Change) < -0.59, p_value < 0.01 are enriched in female hearts. N=2 per sex. (C, D) PCA of data obtained from TMT-MS proteomic characterization in the 8 founder strain hearts. Colors indicate strains and shapes represent sexes, and the variance explained by the two principal components are labeled. (E, F) Correlation of differential proteins across strains in each cellular fraction.

**Figure S4.**
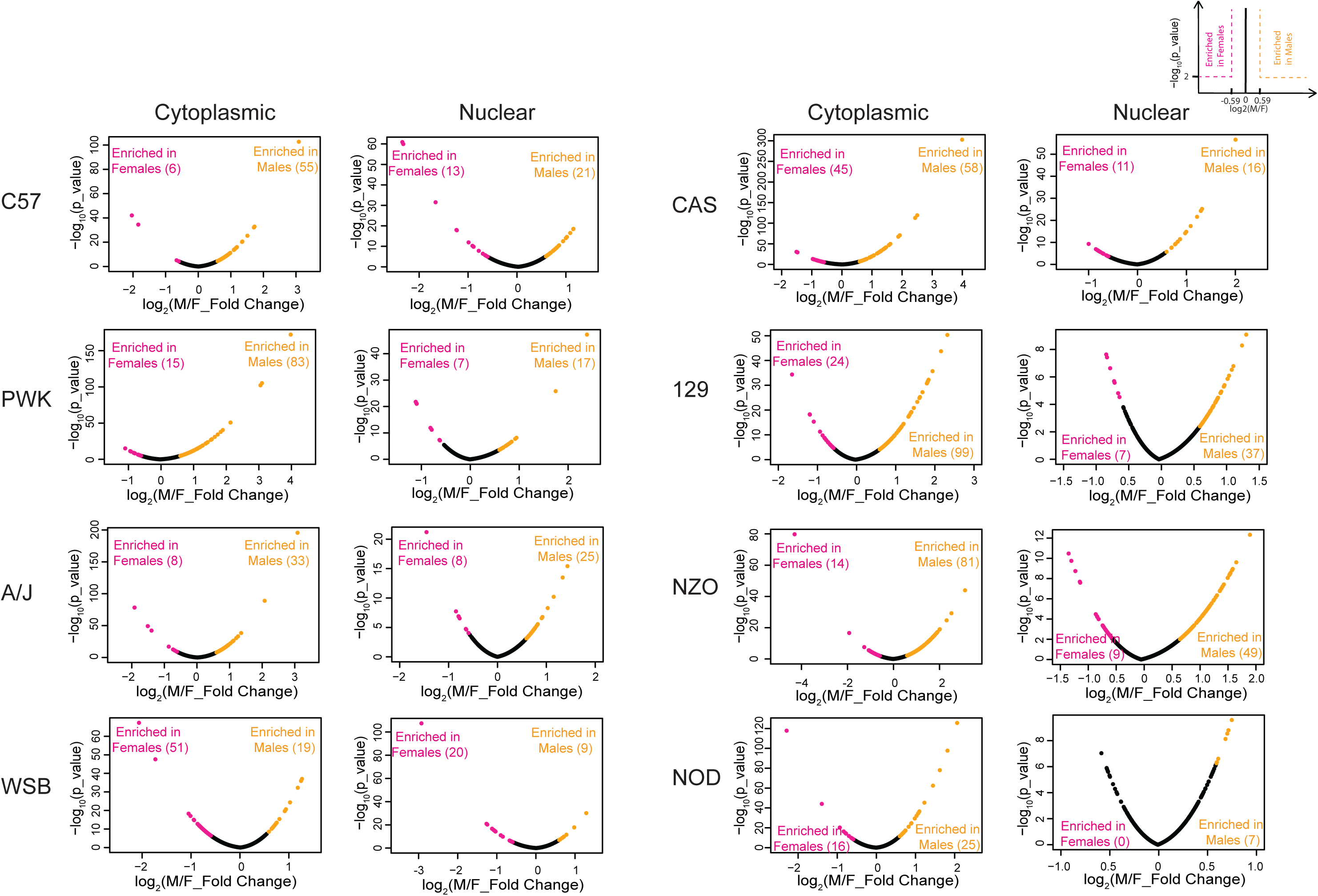
Proteins quantified in each strain. Volcano plots of TMT-MS in each strain in each cellular fraction showing log_2_ (Male/Female_Fold Change) plotted against the –log_10_(p_value). Proteins with log_2_ (Male/Female_Fold Change) > 0.59, p_value < 0.01 are enriched in male hearts; log_2_ (Male/Female_Fold Change) < -0.59, p_value < 0.01 are enriched in female heart. N=2 per sex.

**Figure S5.**
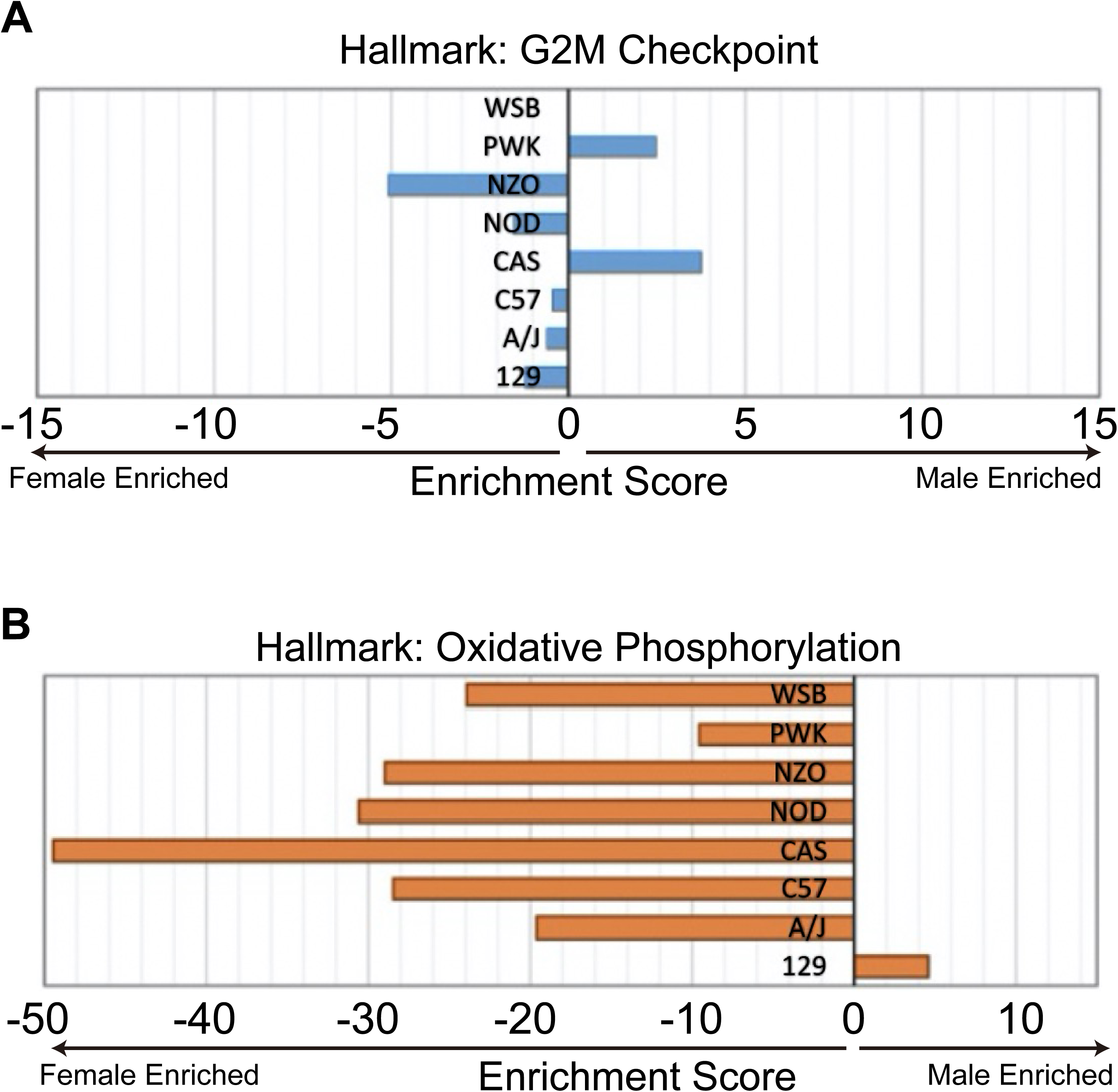
Representative hallmark pathways of each strain. (A) Hallmark enrichment analysis of proteins involved in G2M checkpoint in eight founder strains. (B) Hallmark enrichment analysis of proteins regulating oxidative phosphorylation pathway for founder strains. Degree of enrichment was denoted by Enrichment Score.

**Figure S6.**
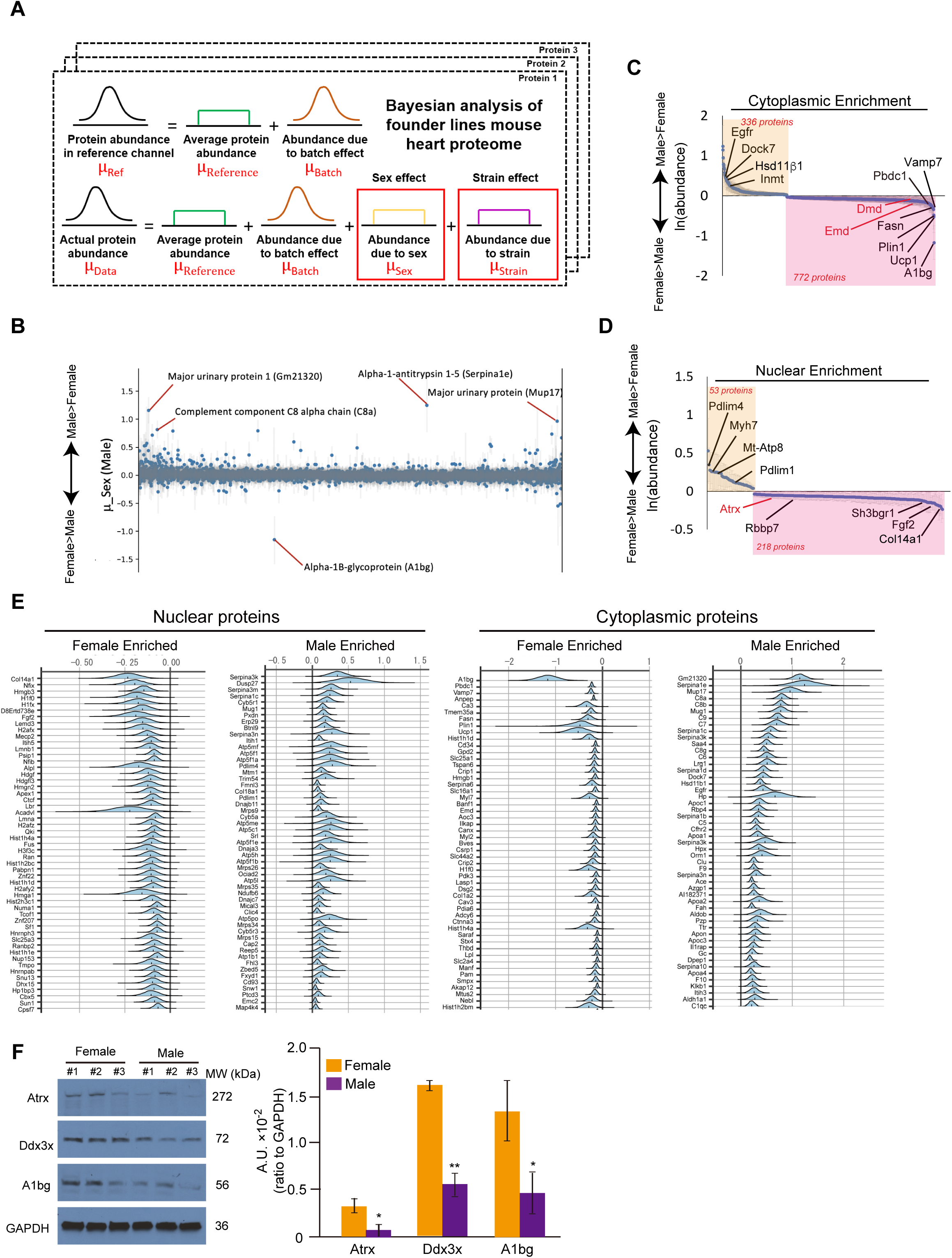
Bayesian inference analysis of quantified proteins in the founder strains. (A) Schematic illustration of Bayesian analysis. Protein abundance of one particular protein in the reference channel is modeled as the addition of the average protein abundance and the batch effect arising from labeling strategies. The protein abundance in each sample is a combination of the reference abundance and abundances ascribed to sex and strain. (B) Sex biases of all quantified proteins inferred by Bayesian analysis. Bars represent mean ± credible interval. (C, D) Waterfall plots of differential protein abundance between male and female hearts in cytoplasm (C) and nuclei (D) based on Bayesian inference analysis. Displayed genes are those that whose 2.5% and 97.5% quantiles of posterior distribution (central 95% Bayesian credible interval) are both greater or both less than 0. The mean of the posterior distribution of each protein and its credible intervals are plotted. Proteins with ln (Male/Female_abundance) > 0 are enriched in males, otherwise enriched in females. Numbers of differential protein are shown in each fraction of both genders, and proteins highlighted in Figure 2c are labeled red. (E) Top 50 significantly differential proteins, based on Bayesian inference, in male and female founder strain hearts identified in nuclei (left) and cytoplasm (right), respectively. (F) Western blot and quantification of protein abundance of Atrx, Ddx3x, and A1bg in male and female hearts. N=3 per sex, *p < 0.05, **p < 0.01, error bars represent mean ± SEM. Data is represented relative to GAPDH levels.

**Figure S7.**
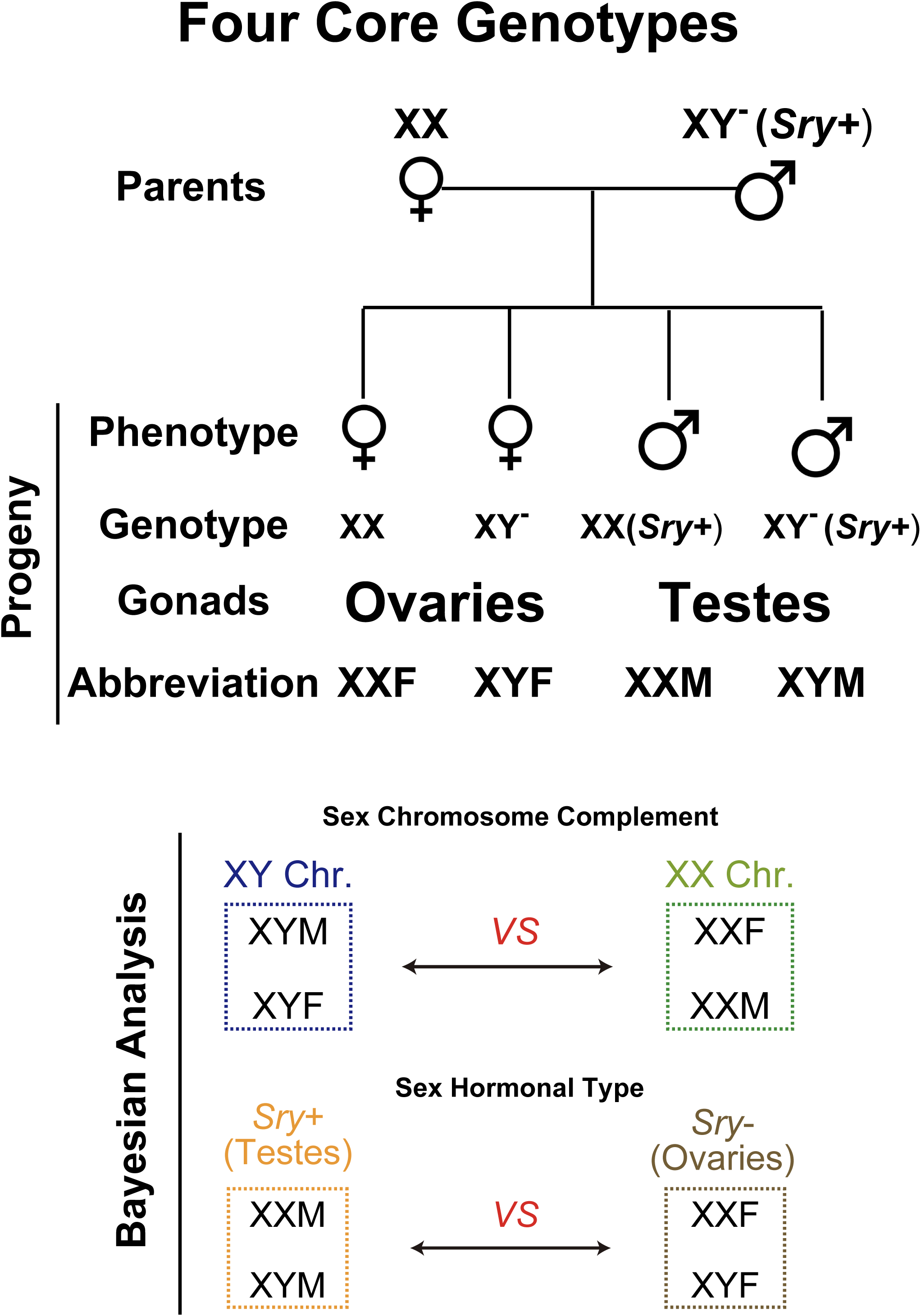
Schematic of the Four Core Genotype (FCG) mouse model. In the FCG model, the testis-determining gene (*Sry*) is removed from Y chromosome to make the “Y^-^” chromosome, and an *Sry* transgene is inserted into chromosome 3. XY^-^ mice have ovaries and called XY females (XYF). Mice with an *Sry* transgene have testes and are called males (XXM or XYM). Mating XX females (XXF) to XY^-^(*Sry*+) (XYM) yields progeny with four genotypes: XXF and XYF (both are females and develop ovaries), XXM and XYM (both are males and develop testes). This model can discriminate between sex differences determined by gonadal type (presence / absence of *Sry*) (XXM and XYM differ from XXF and XYF) versus those determined by the effects of sex chromosomes (XXF and XXM differ from XYF and XYM).

**Figure S8.**
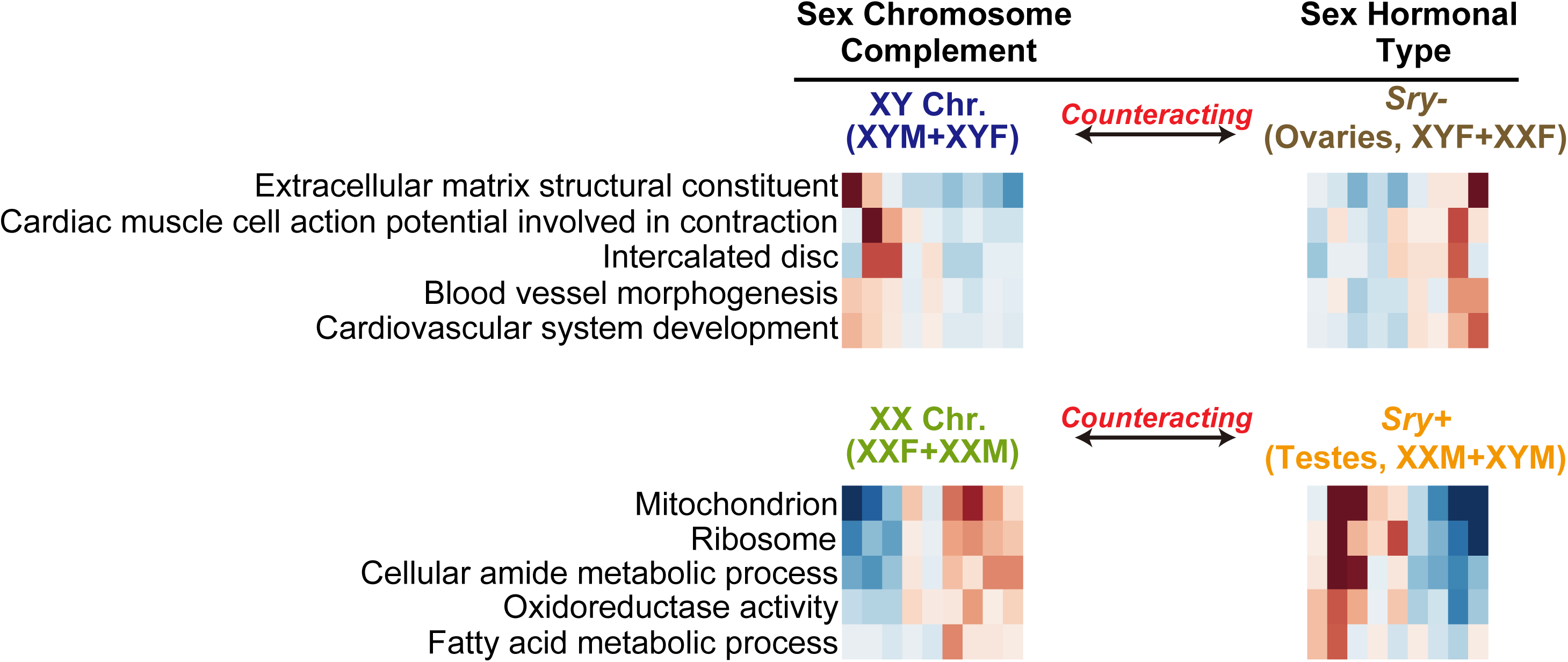
Counteracting effects of sex chromosomes complement and sex hormones. Overlapping enriched GO terms driven by XY chromosomes and ovary hormones (top) and driven by XX chromosomes and testes hormones (bottom). The overlaid biological pathways are regulated by opposing effects of sex chromosomes and sex hormones in normal males (XYM) and females (XXF).

**Tables S1-S10. Lists of differentially expressed genes or proteins in specific experiments or analyses**. (Tables are uploaded as a single Excel spreadsheet (Data file S1).)

Table S1: Differential Genes in Male and Female C57 Mouse Hearts

Table S2: Differential Proteins in Male and Female C57 Mouse Hearts

Table S3: Normalized TMT/MS Data for Nuclear Cardiac Proteins across 8 Founders

Table S4: Normalized TMT/MS Data for Cytoplasmic Cardiac Proteins across 8 Founders

Table S5: Correlation Matrix of Differential Cardiac Proteins across 8 Founders

Table S6: Differential Cardiac Proteins across 8 Founders based on Bayesian Analyses

Table S7: Relative Abundance of all Identified Cardiac Proteins across the Four Core Genotypes

Table S8: Differential Cardiac Proteins across the Four Core Genotypes based on Bayesian Analyses

Table S9: Identified Cardiac Proteins in C57 E9.5 Embryonic Hearts

Table S10: Differential Cardiac Proteins in C57 E9.5 Embryonic Hearts

**Raw images and blots used in figures (Data file S2)**.

Uncropped images of IHC (Figure 2E) and Western blots (Figure S6F).

