## Supplementary material for "Cardiac Sex Differences are Established Prior to Gonad Formation": Key Resource Table

**KEY RESOURCES TABLE**

| REAGENT or RESOURCE | SOURCE | IDENTIFIER |
| --- | --- | --- |
| Antibodies | | |
| Rabbit monoclonal anti-Emerin (Emd) | Abcam | Cat# ab156871 |
| Rabbit polyclonal anti-Dystrophin (Dmd) | Abcam | Cat# ab15277; RRID: AB_301813 |
| Rabbit polyclonal anti-Atrx | Abcam | Cat# ab97508; RRID: AB_10680289 |
| Rabbit polyclonal anti-A1bg | Abcam | Cat# Ab231805 |
| Mouse monoclonal anti-GAPDH | EMD Millipore | Cat# MAB374; RRID: AB_2107445 |
| Mouse monoclonal anti-Tropomyosin | DHSB | Cat# CGbeta6; RRID: AB_10573118 |
| Alexa Fluor 488 goat anti-mouse IgG H+L | Thermo | Cat# A-11001; RRID: AB_2534069 |
| Alexa Fluor 546 goat anti-rabbit IgG1 | Thermo | Cat# A-21123; RRID: AB_2535765 |
| Peroxidase-IgG Fraction Monoclonal Mouse Anti-Rabbit IgG, Light Chain Specific | Jackson ImmunoResearch Labs | Cat# 211-032-171; RRID: AB_2339149 |
| Peroxidase-AffiniPure Donkey Anti-Mouse IgG antibody | Jackson ImmunoResearch Labs | Cat# 715-035-150; RRID: AB_2340770 |
| Bacterial and Virus Strains | | |
| N/A | N/A | N/A |
| Biological Samples | | |
| N/A | N/A | N/A |
| Chemicals, Peptides, and Recombinant Proteins | | |
| DAPI | Thermo | Cat# D1306 |
| Critical Commercial Assays | | |
| RNeasy Plus Mini Kit | QIAGEN | Cat# 74136 |
| TRIzol Reagent | Thermo | Cat# 15596018 |
| RQ1 RNase-Free DNase | Promega | Cat# M6101 |
| BCA Protein Assay Kit | Thermo | Cat# 23225 |
| Tissue-Tek OCT Compound | Sakura | Cat# 4583 |
| Deposited Data | | |
| MS/MS Raw files and MaxQuant analysis files | ProteomeXchange | ProteomeXchange: PXD020139 |
| Experimental Models: Cell Lines | | |
| N/A | N/A | N/A |
| Experimental Models: Organisms/Strains | | |
| Mouse: C57BL/6J | The Jackson Lab | N/A |
| Mouse: PWK/PhJ | The Jackson Lab | N/A |
| Mouse: A/J | The Jackson Lab | N/A |
| Mouse: WSB/EiJ | The Jackson Lab | N/A |
| Mouse: CAST/EiJ | The Jackson Lab | N/A |
| Mouse: 129S1/SvImJ | The Jackson Lab | N/A |
| Mouse: NZO/H1LtJ | The Jackson Lab | N/A |
| Mouse: NOD/ShiLtJ | The Jackson Lab | N/A |
| Mouse: FCG | Arthur Arnold’s Lab | N/A |
| Oligonucleotides | | |
| Primer for genotyping the *Sry* gene, Fwd: 5’- TTGTCTAGAGAGCATGGAGGGCCATGTCAA-3’ | This paper | N/A |
| Primer for genotyping the *Sry* gene, Rev: 5’- CCACTCCTCTGTGACACTTTAGCCCTCCGA-3’ | This paper | N/A |
| Primer for genotyping the control gene (*Fabpi*), Fwd: 5’- CCTCCGGAGAGCAGCGATTAAAAGTGTCAG-3’ | This paper | N/A |
| Primer for genotyping the control gene (*Fabpi*), Rev: 5’- TAGAGCTTTGCCACATCACAGGTCATTCAG-3’ | This paper | N/A |
| Recombinant DNA | | |
| N/A | N/A | N/A |
| Software and Algorithms | | |
| ImageJ (version 1.53a) | National Institutes of Health | https://imagej.net/Downloads |
| R v3.7 | R Project for Statistical Computing | https://www.r-project.org/ |
| Proteome Discoverer 2.3 | Thermo Scientific | N/A |
